# Mast cells and γδ T cells are largely dispensable for adaptive immune responses after laser-mediated epicutaneous immunization

**DOI:** 10.1101/774323

**Authors:** Isabella Joubert, Daniel Kovacs, Sandra Scheiblhofer, Petra Winter, Evgeniia Korotchenko, Helen Strandt, Richard Weiss

**Affiliations:** Department of Biosciences, University of Salzburg, Salzburg, Austria; Department of Hematology and Medical Oncology, Klinikum rechts der Isar and TranslaTUM Cancer Center, Technische Universität München, München, Germany

**Keywords:** epicutaneous vaccination, skin immunology, innate immunity, Phl p 5, glycoconjugates, mannan, barrier disruption

## Abstract

**Background:** The skin resembles an attractive target for vaccination due to its accessibility and abundance of resident immune cells. Cells like γδ T cells and mast cells (MCs) are part of the first line of defence against exogenous threats. Despite being important mediators for eliciting TH2 immune responses after epithelial stress, γδ T cell and MC function still remains to be completely understood. Here, we aimed to characterize their roles in shaping adaptive immune responses after laser-mediated epicutaneous immunization (EPI).

**Methods:** γδ T cell knock out, MC depleted, and wildtype control mice were immunized with mannan-conjugated grass pollen allergen Phl p 5 (P5-MN) by laser-mediated EPI. After 2-3 immunizations, cytokine expression, T helper polarization, and antigen-specific IgG1/IgE levels were analysed. The local cytokine/chemokine milieu after laser microporation was determined.

**Results:** While the majority of inflammatory chemokines and cytokines induced by laser treatment was not affected by the presence of γδ T cells or MCs, RANTES, was elevated in γδ T cell knock out mice, and GROα and TSLP, were significantly decreased after MC depletion. However, absence of γδ T cells or depletion of MC had no substantial effect on adaptive humoral or cellular immune responses after laser-mediated EPI, except for slightly reduced IgG1 and effector T cell levels in MC depleted mice.

**Conclusions:** γδ T cells did not play a pivotal role in shaping the humoral and cellular adaptive immune response after laser-mediated EPI, whereas MC depletion decreased numbers of effector T cells, indicating a potential role of MCs in the activation and maturation of T cells after EPI.

**Highlights:**

- Laser microporation induces an inflammatory chemokine milieu at the site of immunization
- γδ T cells and mast cells contribute to the steady-state or damage-induced cytokine milieu in the skin
- γδ T cells and mast cells are dispensable for adaptive immunity after laser-mediated immunization

## 1. Introduction

While vaccination via intramuscular (IM) or subcutaneous (SC) injection represents a straightforward and well-established method, in most cases, large doses of the respective vaccine are required to provide protective immunity. Moreover, due to the low number of resident antigen-presenting cells (APCs) in these target tissues, IM or SC inoculations require infiltrating leukocytes attracted by inflammatory signals at the site of vaccine administration [1]. To overcome the limitations of conventional injections, new methods have recently been developed with the focus on generating less invasive and more efficient delivery techniques for vaccines [2, 3].

Whereas some novel skin vaccination technologies rely on actual deposition of the vaccine within the skin, others aim at passive transport of the vaccine via barrier disrupted skin, e.g. by using laser microporation [3, 4]. Thereby, heat pulses are produced by highly focused thermal energy (e.g. infrared light) leading to thermal ablation at locally restricted skin areas [5] and thus generating skin micropores in a controlled manner by means of decomposition and vaporization of tissue. Laser microporation enables circumvention of the stratum corneum and promotes vaccine delivery to cutaneous APCs. In addition to enhanced antigen delivery, this technique provides an intrinsic adjuvant effect [6] and has shown promising results in prophylactic and therapeutic approaches against type I allergies and tumours in animal models [4, 7, 8].

As the barrier between host and environment, the skin is a highly immunocompetent organ harbouring diverse populations of resident immune cells including dendritic cells (DC), mast cells (MC), and γδ T cells, and thus represents an attractive target for vaccination [9]. While skin-resident APCs are continuously migrating to regional lymph nodes and the blood, sessile (epi)dermal cells are able to polarize immune responses by release of cytokines. An accumulating body of evidence suggests that vaccination via the skin enables dose sparing [10, 11], which is of special interest for treatment of immune-compromised and elderly patients as well as in cases of limited vaccine availability. The easily accessible upper layers of the skin furthermore allow for usage of smaller needles or even needle-free vaccination. This results in increased patient convenience and compliance and reduces the risk of needle-stick injuries, thereby also offering advantages for healthcare providers [12].

In the context of epicutaneous immunization (EPI), antigens are delivered to APCs mostly via passive uptake. Based on the low efficiency of passive uptake, novel antigen formulations directly target receptors on skin-resident DCs to promote antigen uptake. One approach is to actively target DCs by engagement of their expressed C-type lectin receptors (CLRs) and thus significantly increase antigen uptake [13]. Not only does targeting CLRs increase antigen uptake and enhance immunogenicity, but also allows shaping of the subsequent immune responses [14]. CLRs comprise a large superfamily of receptors, which recognize diverse carbohydrate moieties in a calcium-dependent or independent manner through one or more conserved carbohydrate recognition domains (CRDs) or C-type lectin domains (CTLDs). CLRs have evolved alongside pathogens and are often specific for distinct and conserved carbohydrate motifs derived from viruses, bacteria, parasites or fungi [15]. A variety of immune cells express CLRs such as dermal DCs (dDCs), Langerhans cells (LCs), macrophages, neutrophils, and B lymphocytes. As pattern recognition receptors (PRRs), CLRs play various roles in eliciting immune responses depending on the detected pathogen-associated molecular pattern (PAMP). The potential of the carbohydrate mannan as an adjuvant capable of enhancing and shaping the overall immune response has been demonstrated to rely on its capacity to engage CLRs [16, 17]. We have previously reported that EPI with allergen-mannan neoglycoconjuates by laser microporation leads to more efficient antigen uptake by dDCs and LCs. In addition, we found that the initiated immune response was skewed towards TH1/TH17, which is in accordance with the fungal origin of mannan [17].

Nevertheless, the immune response mounted upon an antigen encounter is also highly dependent on the microenvironment within the target tissue. Barrier disruption has been demonstrated to be essential for enhancing skin permeation by altering the microarchitecture of the skin and increasing the motility of embedded APCs. Furthermore, disrupting the stratum corneum has been shown to provide a natural inherent adjuvant effect by inducing the secretion of pro-inflammatory cytokines, chemokines and danger-associated molecular patterns (DAMPs) [18, 19]. More precisely, damaged and necrotic keratinocytes release cytokines such as IL-25, IL-33 and TSLP (“alarmins”), which activate other skin-resident cells and induce specific immune responses [20].

MCs and γδ T cells are distributed throughout the whole body in the mucosal, epithelial and connective tissues. Hence, they are strategically positioned at the barrier between host and environment, where exogenous material can enter. They both play important roles in innate immunity and can be rapidly activated by stress signals and pathogens. The various protective functions exerted by γδ T cells include anti-microbial defence, tumour surveillance, tissue homeostasis, and wound repair [21, 22]. The term lymphoid stress-surveillance is commonly used to describe the rapidly induced immune response initiated after stress-antigen engagement by the γδ T cell receptor. Additionally, γδ T cells have been reported to directly sense epithelial dysregulation, control tissue homeostasis by IL-13 production and promote IgE-dependent responses to protein allergens in stressed tissue [23]. MCs are predominantly known for their essential role as mediators of type I allergy and for their potential to secrete a variety of allergic mediators upon activation [24]. However, MCs cannot only be activated by antigen-dependent crosslinking of IgE:FcεRI, but also by a variety of other stimuli, such as cytokines, neuropeptides and danger signals [25, 26]. Thus, MCs can - depending on the context - elicit pro-inflammatory, anti-inflammatory as well as immunoregulatory responses by releasing distinct patterns of mediators and cytokines [24], [25].

In addition to their protective functions and their role in allergic diseases, both γδ T cells as well as MCs, are implicated in autoimmunity and malignancy, although the nature and extent of their involvement in the etiopathology is debated [27, 28].

In the present study, we investigated the role of γδ T cells and MCs in shaping the adaptive immune response following EPI using mannan-conjugated grass pollen allergen Phl p 5 after laser microporation of skin.

## 2. Material and Methods

### Generation of allergen-mannan neoglycoconjugates

Allergen-mannan neoglycoconjugates were generated by mild oxidation of the mannan moiety and subsequent reductive amination with recombinant Phl p 5 (rPhlp 5) as previously described [17], using a ratio of 5:1 (w/w) between carbohydrate and allergen. For more details, see supplementary methods.

### Purification and characterization of Phl p 5-mannan neoglycoconjugates

Phl p 5 mannan neoglycoconjugates (P5-MN) were subjected to size exclusion chromatography using a 16/60 Sephacryl S-300 HR column (GE Healthcare) (Suppl. Fig. 1). Coupling efficiency of individual glycoconjugate fractions was monitored by a 12.5% reducing SDS-PAGE using colloidal Coomassie staining for protein visualization after which high molecular weight fractions were pooled (Suppl. Fig. 2). The hydrodynamic radius of this glycoconjugate pool was determined using dynamic light scattering (DLS). Measurements were performed at a protein concentration of 1 mg/mL in PBS at 40% laser power for 20×5 seconds on a Viscotec DLS 802 dynamic light scattering instrument (Suppl. Fig. 3).

### Animal experiments

#### Mice

Tcrdt^m1Mom^ targeted mutant strain mice deficient in receptor expression in all adult lymphoid and epithelial organs were purchased from the Jackson Laboratory. Mice were backcrossed for 14 generations to a BALB/c genetic background. For all experiments, homozygous T-cell receptor delta chain (TCRd) knock-out mice and heterozygous littermates (controls) were used.

Transgenic Mas-TRECK (mast cell-specific enhancer-mediated toxin receptor-mediated conditional cell knock out) mice have been previously described by Otsuka et al. and Sawaguchi et al. [29, 30]. MCs of this transgenic strain express the human diphtheria toxin receptor (hDTR) under the control of an intronic enhancer (IE) element of the IL-4 gene. MCs in Mas-TRECK mice were conditionally depleted by intraperitoneal injection of 250 ng Diphtheria toxin (DT) for five consecutive days prior to the first immunization. Treatment with DT also leads to the complete temporary depletion of basophils, since they express low levels of hDTR. C57BL/6 mice obtained from Charles River Laboratories, Sulzfeld, Germany were used as controls. Prior to the experiments, all mice were genotyped via PCR. To ensure the absence of MCs and γδ T cells at the time of immunization, epidermal sheets and histological cryosections were prepared (Suppl. Fig. 4 and 5).

TCRd, Mas-TRECK and C57BL/6 mice were kept in the animal facility of the University of Salzburg and were maintained according to the local animal care guidelines. All animal experiments were approved by the Austrian Ministry of Science (permit number BMWFW-66.012/0014-WF/V/3b/2017).

#### Immunizations

Immunizations were performed by laser microporation of the skin using the P.L.E.A.S.E© laser device (Erbium-Yttrium aluminium garnet laser, Pantec biosolutions) followed by application of P5-MN in solution to the laser-generated micropores as previously described [4]. Laser microporation was performed with the following settings: 500 Hz, pulse duration of 50 µs, 5% pore density, 3 pulses per pore and a total fluence of 8.4 J/cm^2^. TCRd ^−/−^ and heterozygous control mice were immunized on days 0, 14 and 28 with 1 µg P5-MN. MC-depleted Mas-TRECK and C57BL/6 control mice were immunized on days 0 and 14 with 1 µg Phl p 5 mannan.

### Antigen-specific antibody measurements

10-13 days after the last immunization, mice were sacrificed, and blood samples were drawn either from the saphenous vein or the retrobulbar sinus. Phl p 5-specific IgG1 antibodies were detected by luminometric ELISA. To assess cell-bound Phl p 5-specific IgE, a basophil activation test was conducted as described [31, 32].

### Lymphocyte cultures, flow cytometry, and cytokine measurements

Isolated lymphocytes from spleens and lymph nodes were cultured in a sterile 96-well tissue culture plate (U-bottom; Greiner) at a density of 4×10^6^ cells/mL in the presence of 10 µg/mL rPhl p 5 in T cell medium (RPMI-1640; 10% FCS, 25mM HEPES, 2mM L-Glu, 100 µg/mL streptomycin, 100 U/mL penicillin) for five days. At day five, culture supernatants were harvested, and cytokine profiles were assessed either by using ProcartaPlex multiplex cytokine panels (Thermo Fisher) or LegendPlex Mouse Th cytokine panel (BioLegend), respectively. On the same day, restimulated cells were analysed by flow cytometry. In brief, cells were washed with DPBS, blocked with an anti-CD16/32 containing hybridoma supernatant, and stained for the extracellular markers using anti-CD4 (APC-Cy7-labelled; clone GK1.5, eBioscience; 1:400), anti-CD44 (BV650-labelled; clone IM7, BioLegend; 1:100), anti-CD62L (FITC-labelled; clone MEL-14, BioLegend; 1:200) and live/dead staining (fixable viability dye eFluor 506; Thermo Fisher; 1:1000) for 30 min on 4°C. For intracellular staining, cell fixation and permeabilization was performed using the FoxP3 Staining Buffer Sets (Tonbo Biosciences) according to the manufacturer’s instructions and 10% naïve mouse serum (in permeabilization buffer) was used for additional blocking. After 30 min of incubation with intracellular markers for FoxP3 (APC-labelled; clone FJK-16s, Invitrogen; 1:100), GATA3 (BV421-labelled; clone 16E10 A23, BioLegend; 1:200), RORγT (PE-labelled; clone B2D, Thermo Fisher; 1:100) and T-bet (PE-Cy7-labelled; clone 4B10, BioLegend; 1:100) at RT in the dark, cells were washed twice with permeabilization buffer and dissolved in FACS buffer (PBS, 1% BSA, 2mM EDTA) for analysis using a CytoFLEX S flow cytometer (Beckman Coulter).

### Laser-induced cytokine-milieu in murine skin

Skin areas of TCRd^−/−^, TCRd^−/+^, DT-treated and untreated Mas-TRECK mice were excised 6 hours and 24 hours after laserporation and then homogenized in 20-fold volume of DPBS containing 1% of a protease inhibitor cocktail (Sigma, P8340) using an Ultra-Turrax homogenizer (Ika, Staufen, Germany). Samples were centrifuged for 10 min at 21.000 x g, and then the supernatants were 0.22 µm filtered and finally analysed for cytokine/chemokine release using the ProcartaPlex multiplex assay (Thermo Fisher)

### Statistical analysis

Unpaired or paired Student’s t-tests were used for comparing two groups. Effect of genotype and laser treatment on skin chemokines/cytokines was assessed by two-way ANOVA followed by Tukey’s or Holm-Sidak’s post hoc test as indicated (Prism 6, GraphPad Software). Data are expressed as means ± SEM. (P-value range is indicated: *P<0.05, **P<0.01 and ***P<0.001). Flow cytometry profiles and microscopy images are representative of repeated experiments.

## 3. Results

### Contribution of Mast cells and γδ T cells to the cytokine/chemokine milieu at the site of laser microporation

Knowing that both γδ T cells and MCs are sentinels and damage sensors, we evaluated the contribution of these cell types to the locally-induced cytokine and chemokine milieu after laser-induced skin damage. To this end, the skin of TCRd^−/−^ TCRd^−/+^, DT-treated and untreated Mas-TRECK mice was analysed for cytokine/chemokine secretion either 6 h or 24 h after laser microporation. Non-laser microporated mice were included as controls. As expected, in wild-type mice, laser microporation led to an increase of a broad panel of cytokines and chemokines. Both in TCRd^−/−^, TCRd^−/+^ and Mas-TRECK mice, cytokines GROα, MCP-1, TSLP, IL-6, TNF-α, and MCP-5 peaked 6 h after laser microporation (Fig. 1, A and B), whereas the highest levels of MIP-1α, MIP-2, MIP-3α, RANTES, and IP-10 were observed at the 24 h time point. Contrary, MIP-1α, MIP-2, MIP-3α, RANTES, and IP-10 peaked earlier (6 h) in Mas-TRECK mice (Fig. 1B). Tables 1 and 2 summarize the significant changes in cytokine/chemokine concentrations in the skin following laser microporation both in TCRd^−/−^, TCRd^−/+^and Mas-TRECK mice. γδ T cells did not alter the locally-induced cytokine/chemokine milieu after laser microporation, with the exception of RANTES, which was significantly higher in TCR^−/−^ mice. Also, steady-state IL-6 (NS) and MCP-5 (P<0.05) levels were decreased in absence of γδ T cells (Fig. 4A). In accordance with that, we found that IL-6 and MCP-5 peak levels after laser microporation were elevated in the presence of γδ T cells, confirming a significant contribution of γδ T cells to the production/regulation of these chemokines (also see Table 1).

**Figure 1:**
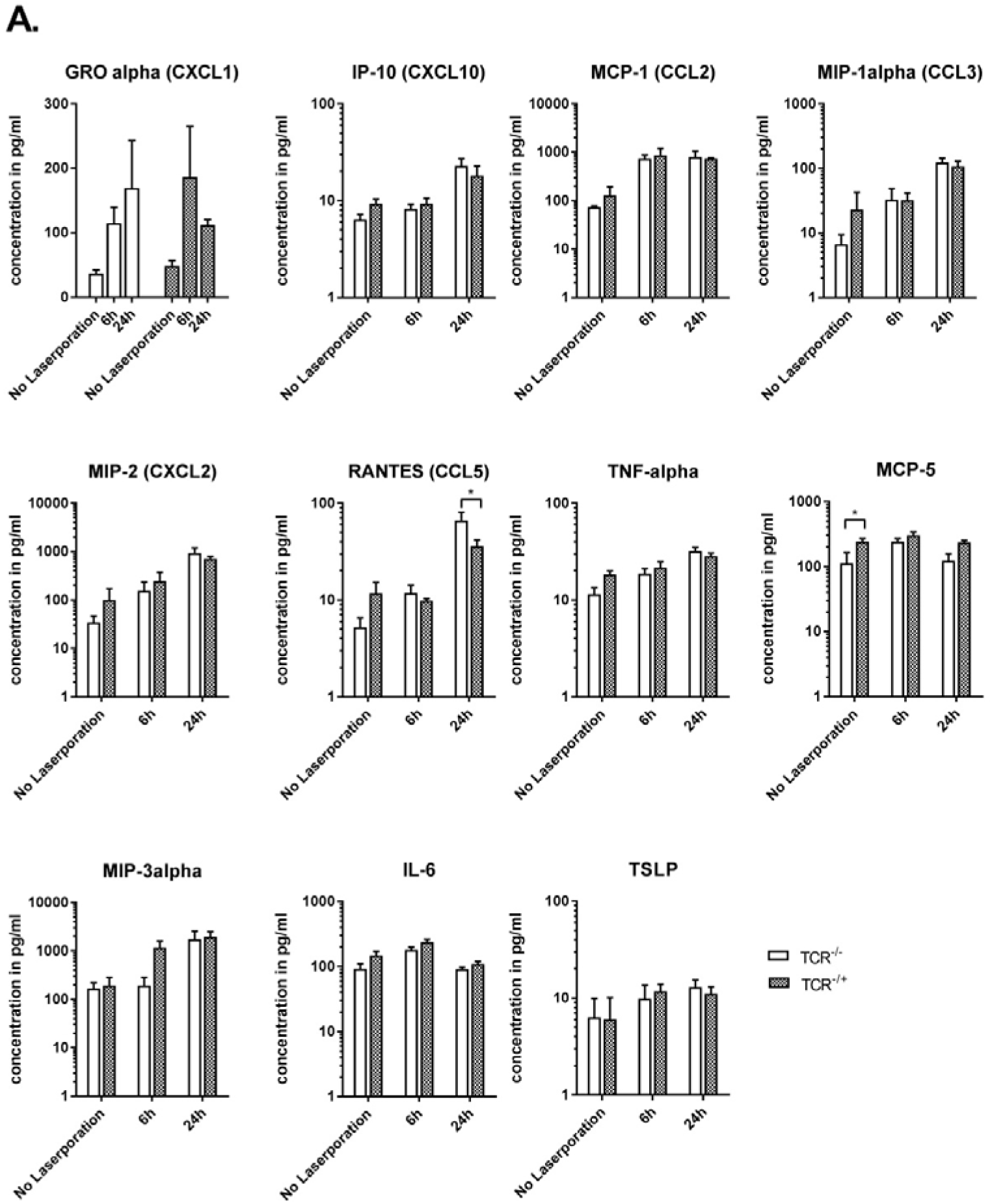
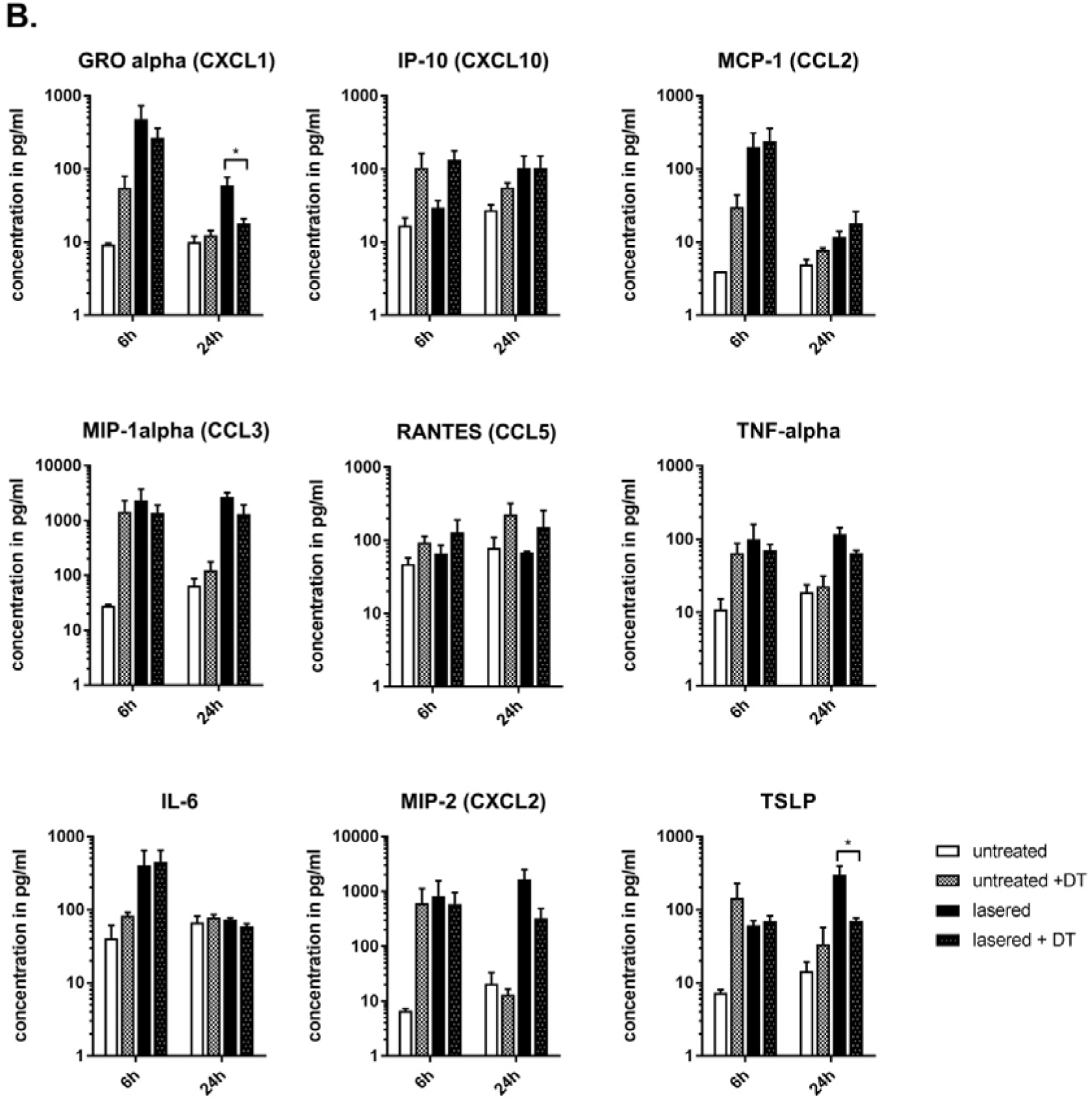
Cytokine/chemokine milieu at the site of laser microporation. Cytokine/chemokine concentrations measured in supernatants of homogenized skin from TCRd (A) or MasTRECK (B) mice 6 h and 24 h after laser microporation using cytokine multiplex assay. Non-laser microporated mice (untreated) were used as controls. Data (n=4) is depicted as cytokine/chemokine concentration in pg/mL. Statistical significance between TCRd^−/−^ and TCRd^+/−^ mice or MC depleted (+DT) vs non-depleted Mas-TRECK mice was assessed by two-way ANOVA followed by Holm-Sidak’s multiple comparison post hoc test. * P<0.05.

**Table 1:** Laser-related effects on the cytokine/chemokine milieu 6 h or 24 h in TCRd mice after laser microporation. Data was analysed by two-way ANOVA (time 0h, 6h, 24h vs. genotype) and Tukey’s multiple comparisons test (0h vs. 6h and 0h vs. 24h). * P<0.05, ** P<0.01, *** P<0.001, **** P<0.0001, NS = not significant.

| Cytokine/Chemokine | 6h laser effect | 24h laser effect | genotype effect |
| --- | --- | --- | --- |
| GROα | NS | NS | NS |
| IL-6 | *** | NS | * |
| IP-10 (CXCL10) | NS | ** | NS |
| MCP-1 (CCL2) | NS | ** | NS |
| MIP-1α (CCL3) | NS | **** | NS |
| MIP-2 (CXCL2) | NS | **** | NS |
| RANTES (CCL5) | NS | **** | NS |
| MIP-3α | NS | ** | NS |
| TNF-α | NS | *** | NS |
| TSLP | NS | NS | NS |
| MCP-5 | * | NS | * |

**Table 2:** Laser- and DT-related effects on different skin cytokines/chemokines 6 h or 24 h after laser microporation in MasTRECK mice were analysed by two-way ANOVA (DT treatment vs. laser treatment). * P<0.05, ** P<0.01, NS = not significant.

| Cytokine/Chemokine | 6h laser effect | 6h DT effect | 24h laser effect | 24h DT effect |
| --- | --- | --- | --- | --- |
| GROα | * | NS | * | NS |
| IL-6 | * | NS | NS | NS |
| IP-10 (CXCL10) | NS | * | NS | NS |
| MCP-1 (CCL2) | * | NS | NS | NS |
| MIP-1α (CCL3) | NS | NS | ** | NS |
| MIP-2 (CXCL2) | NS | NS | * | NS |
| RANTES (CCL5) | NS | NS | NS | NS |
| TNF-α | NS | NS | ** | NS |
| TSLP | NS | NS | * | NS |

In Mas-TRECK mice, we observed a decrease in the secretion of cytokines such as GROα (P<0.05), MIP-1α (NS), MIP-2α (NS), and TSLP (P<0.05) dependent on the presence of MCs after laser microporation (Fig. 1B). In addition, we found a DT-mediated effect on the release of all assessed cytokines/chemokines with the exception of IL-6 and RANTES (significant only for IP-10, see Table 2).

### Mast cells and γδ T cells play no essential role in induction of humoral Th2 responses after EPI

To investigate the role of γδ T cells and MCs in the systemic immune response after EPI with P5-MN, specific IgG1 and cell-bound IgE levels were measured by ELISA and BAT, respectively. Elevated IgG1 serum levels and cell-bound Phl p 5-specific IgE were found in all groups after EPI, suggesting an ongoing humoral immune response. In general, TCRd mice produced higher levels of IgG1 and IgE compared to Mas-TRECK mice (C57BL/6) due to their BALB/c genetic background. Unlike TCRd^−/−^, TCRd^−/+^ mice showed no reduction of antigen-specific IgG1 production after EPI with P5-MN (Fig. 2A). However, we found a significant decrease in serum IgG1 levels in MC-depleted Mas-TRECK mice compared to C57BL/6 WT mice, thereby indicating at least a partial role of MCs in the TH2-dependent switch to IgG1. Despite the differences in IgG1 levels in MC-depleted mice, in our experimental setting, we observed no significant differences in cell-bound antigen-specific IgE. Furthermore, cell-bound antigen-specific IgE levels were comparable between TCRd^−/−^ and TCRd^−/+^ mice (Fig 2B). Thus, both γδ T cells as well as MCs play no major role in the induction of humoral TH2 responses after two (MCs) or three (γδ T cells) epicutaneous immunizations with P5-MN.

### TH2 cytokine production following EPI is elevated in the absence of mast cells

Having found that both γδ T cells and MCs had little effect on humoral TH2 responses, we were prompted to investigate whether these cells influence TH2 cytokine release. To address this question, splenocytes from immunized mice and naïve controls were restimulated with Phl p 5 in vitro for five days and cytokine secretion was measured by ProcartpaPlex/LegendPlex multiplex analysis. Whereas splenocytes from naïve mice displayed only low baseline secretion of cytokines upon Phl p 5 exposure (<10pg/mL for TNF-α, GROα, IL-5, and IL-13; all others <1pg/mL), splenocytes from immunized mice produced significant amounts of pro-inflammatory, TH1, TH2, and TH17 cytokines. As expected, splenocytes from MasTRECK mice secreted higher levels of the TH1 cytokine IFN-γ and lower levels of TH2 cytokines IL-4, IL-5, and IL-13 compared to TCRd mice due to their genetic background. Again, we observed no significant changes in the secretion of cytokines between TCRd^−/−^ and TCRd^−/+^ mice in our experiments (Fig. 3A). We also found no major differences between MC-depleted mice and C57BL/6 control mice except for slightly higher concentrations of IL-4 and IL-10 (Fig. 3B). Additionally, we detected increased levels of the TH2 cytokines IL-4, IL-5 and IL-10 in ex vivo restimulated cells from draining lymph nodes from MC-depleted Mas-TRECK mice (Suppl. Fig. 6). Taken together, these findings suggest that in our experimental setting, the induction of TH2 cytokines was elevated in the absence of MCs, which is in contrast to the observed reduction of IgG1. The secretion of pro-inflammatory, TH1 and TH17 cytokines, however, was found to be independent from the presence of MCs. γδ T cells had no effect on the TH cell profile induced by EPI.

**Figure 2.**
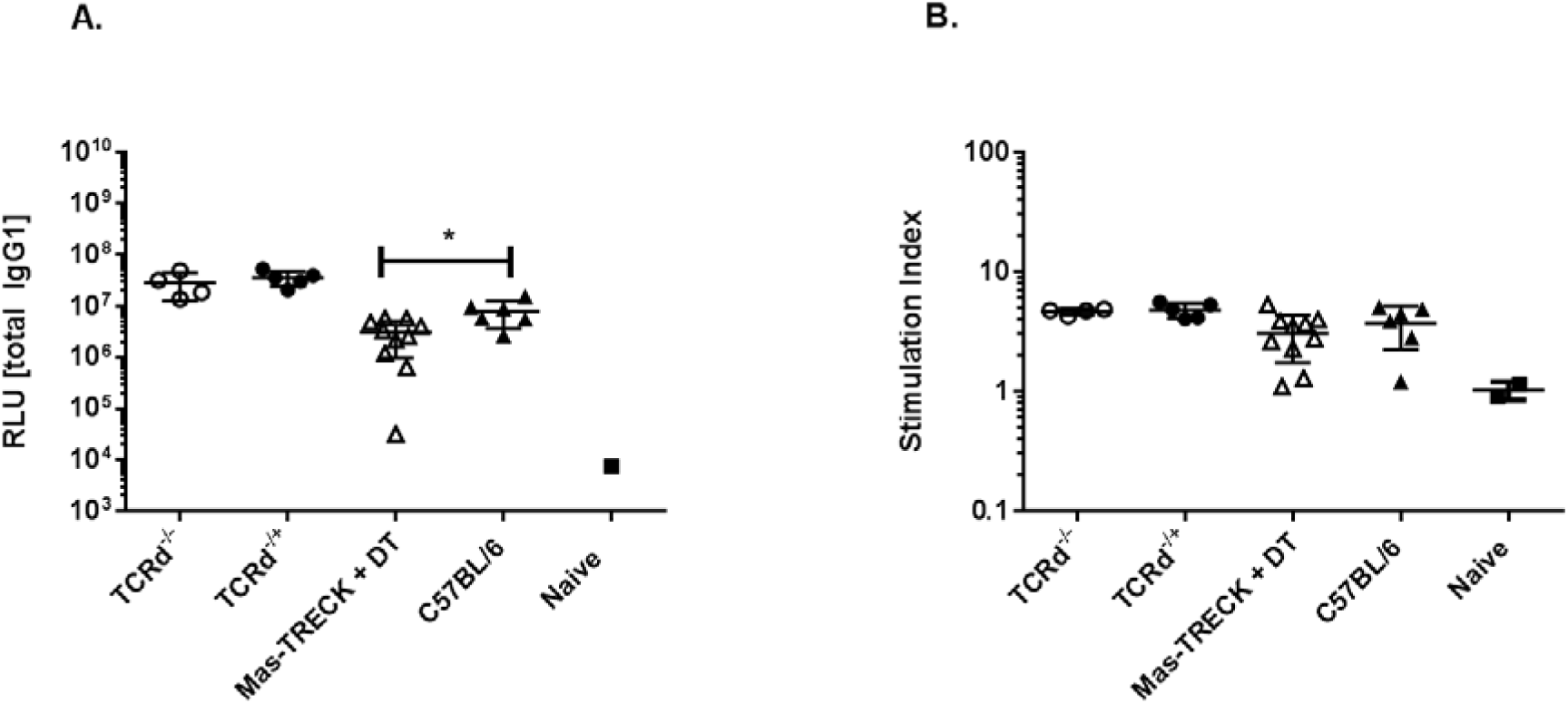
Antigen-specific IgG1 and IgE antibody titers after laser-mediated EPI with P5-MN. **A)** A luminometric ELISA was performed with 1:1000 diluted serum samples and Phl p 5-specific IgG1 levels were measured in terms of RLU (relative light units) **B)** The basophil activation due to IgE crosslinking after restimulation with 10 ng/mL Phl p 5 was measured in terms of CD200R upregulation via flow cytometry. Mean fluorescence intensity (MFI) of CD200R was normalized to untreated samples. TCRd^−/−^ (n=4), TCRd^−/+^ (n=6), DT-treated Mas-TRECK (n=10), C57BL/6 (n=6) mice and naїve (n=1-2) mice. *P<0.05

**Figure 3:**
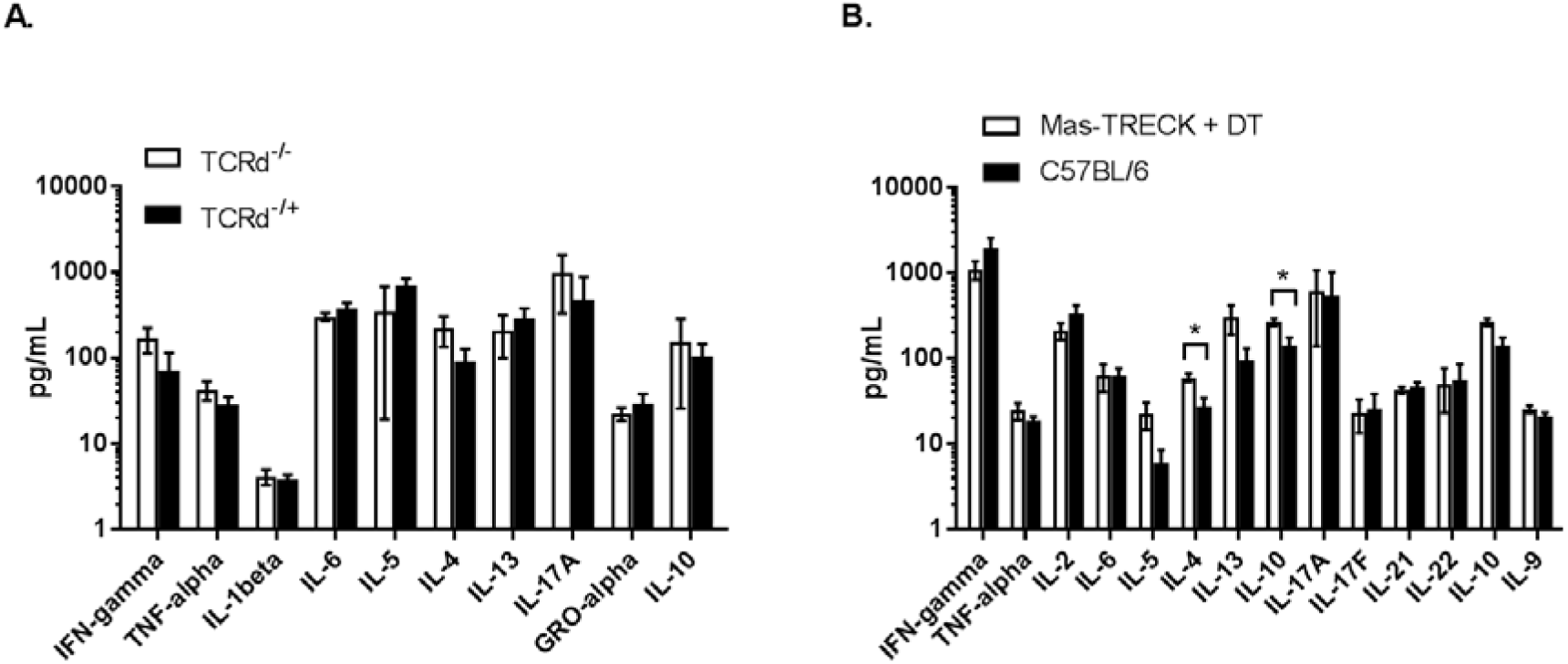
Cytokine profiles of cultured splenocytes from immunized mice. Cytokine production in cell culture supernatants of restimulated splenocytes was measured using ProcartpaPlex (A) or LegendPlex (B) multiplex analysis. **A.)** TCRd^−/−^ (n=4), TCRd^−/+^ (n=5), and **B.)** DT-treated Mas-TRECK (n=10) and C57BL/6 (n=6) mice). *P<0.05

### Presence of mast cells contributes to T cell activation and maturation but not to T helper polarization

Based on the differences we detected in cytokine secretion profiles of restimulated splenocytes and SDLN cells at least in MC-depleted mice, we asked whether γδ T cells or MCs influence T cell activation or polarization.

Analysis of intra- and extracellular staining of Phl p 5-restimulated SDLN cells revealed no differences in the percentages of naïve, central memory, and effector CD4^+^ T cells or the amount of transcription factor expression between TCRd^−/−^ and TCRd^−/+^ groups (Fig. 4). While the percentage of naïve CD4^+^ T cells was significantly higher in MC-depleted mice, the amount of effector T cells was lower, thereby implying a role of MCs in T cell activation and maturation. In particular, we found that expression of key TH transcription factors T-bet, GATA3, RORγt and FoxP3 was independent of MC depletion. Thus, we could not confirm an influence of MCs on TH2 polarization on the transcription factor level.

**Figure 4:**
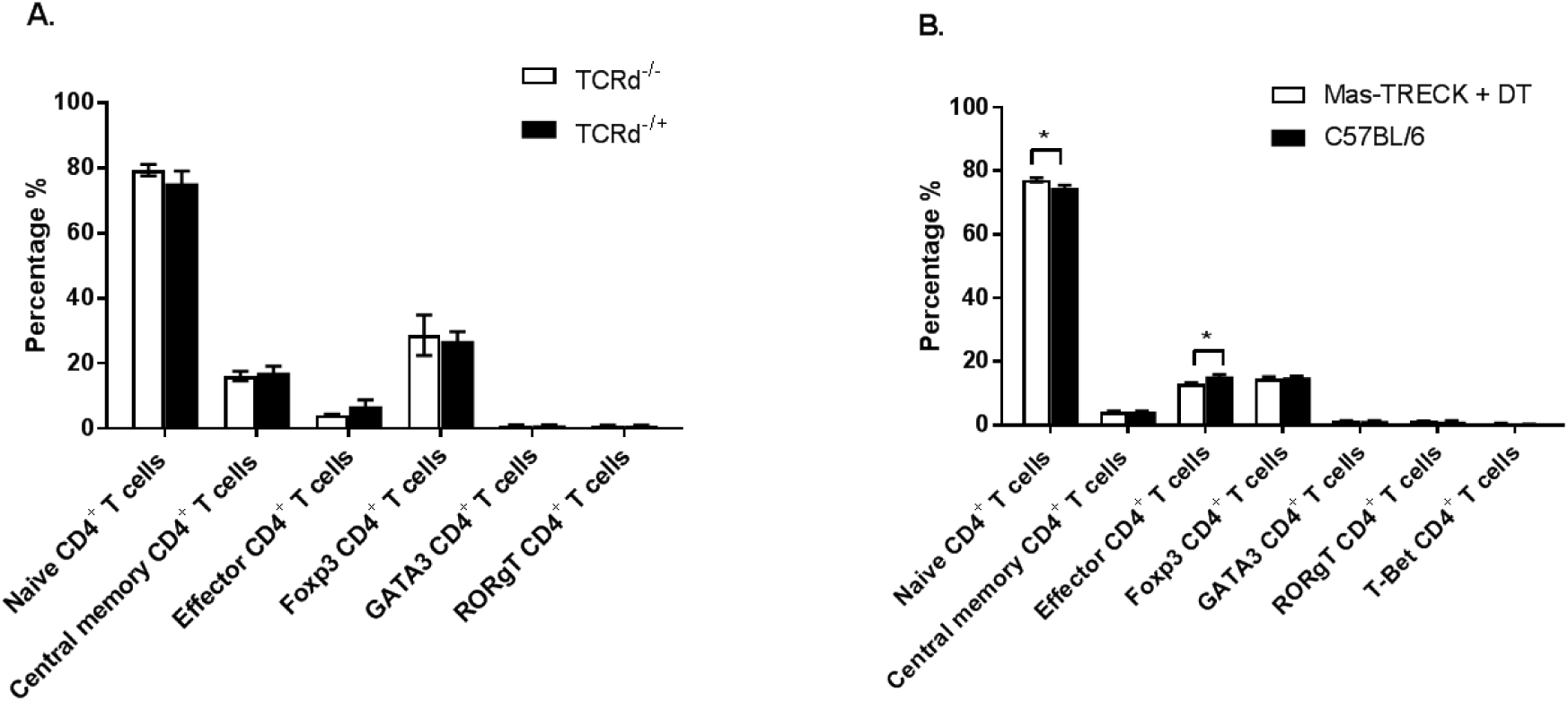
T helper polarization in Phl p 5-restimulated SDLN cells. Extracellular markers CD4, CD62L and CD44 were used to determine the distribution of naïve, central memory, and effector CD4^+^ T cells. Viable and singlet CD4^+^ T cell subset distribution and the percentage of FoxP3, GATA3, RORγT and T-bet-producing CD4^+^ T cells was determined using flow cytometry. **A.)** TCRd^−/−^ (n=4), TCRd^−/+^ (n=5), **B.)** DT-treated Mas-TRECK (n=10) and C57BL/6 (n=6). *P<0.05

## 4. Discussion

Laser microporation represents a novel method for skin-based vaccination exploiting the richness of superficial skin layers in antigen presenting cells. Furthermore, the microporation process itself has been shown to locally create an inflammatory milieu, thereby providing an adjuvant effect, which potentially obviates the need for adjuvants in vaccine formulations. By targeting receptors on dendritic cells with antigen coupled to glycans such as mannan, we have also demonstrated that a synergistic effect between microporation and directly addressing antigen presenting cells can be achieved, which could not be observed following intradermal injection [17]. Whereas the role of various skin-resident dendritic cell types and Langerhans cells in eliciting immune responses following laser-facilitated skin immunization has been studied in detail [33-35], it remains to be elucidated whether innate immune cells contribute to these reactions. In the current study we therefore investigated the impact of γδ T cells and MCs for production of specific antibodies as well as local and systemic chemokine and cytokine secretion following laser-mediated EPI. As antigen we chose the grass-pollen allergen Phl p 5 coupled to mannan, which is a promising candidate for allergen-specific immunotherapy via the skin as it combines high immunogenicity with low allergenicity [17].

Skin-resident dendritic epidermal T cells (DETCs) in mice as well as their human counterparts exclusively express γδ T cell receptors and are able to modulate immune responses initiated in the epithelium. Both have been implicated in maintenance of the immunologic integrity of the skin acting as sensors of cellular dysregulation by recognizing microbial moieties or ligands expressed by tumor cells [23, 36]. As it has been previously shown that DETCs are essential for strong TH2-associated atopic responses upon antigen encounter in context with cutaneous epithelial stress, it was obvious to investigate the role of these cells in immune reactions following antigen delivery via barrier disrupted skin.

Our current study shows that the role of γδ T cells in shaping the local cytokine/chemokine milieu following laser microporation is marginal and they consequently have no effect on eliciting adaptive immune responses after laser-mediated EPI.

Upon activation, DETCs have been reported to be potent producers of pro-inflammatory cytokines such as IL-2, IL-3, GM-CSF, IFN-γ and TNF-α and they can also produce TH1 effector cytokines (IFN-γ, TNF-α), TH2 effector cytokines (IL-4, IL-5, IL-10, IL-13) and TH17 effector cytokines (IL-17) [37, 38]. Additionally, activated γδ T cells are known to secrete the chemokines lymphotactin (XCL1) and RANTES (CCL5) [22] and also produce the macrophage-homing chemokines MIP-1α (CCL3) and MIP-1β (CCL4) [39]. However, we found that the cytokine and chemokine expression in the skin after laser microporation is largely independent of γδ T cell presence (Fig. 1A and Table 1). A surprising finding was that the production of both IL-6 and MCP-5 (CCL12) was reduced in untreated mice lacking γδ T cells, thereby suggesting them to play a role in steady-state production of these chemokines (Fig. 1A).

In our experimental setting, the induction of antigen-specific IgG1 and IgE (Fig 1.) and the secretion of TH2 cytokines such as IL-4, IL-5, IL-10 and IL-13 of *in vitro* restimulated splenocytes (Fig. 3A) do not differ between TCRd^−/−^ and TCRd^−/+^ mice and therefore do not depend on the presence of γδ T cells.

In contrast, Strid et al. have demonstrated a crucial role for γδ T cells in producing antigen-specific IgG1 and IgE after allergen application to stressed epithelia [23]. However, in this study a different method for barrier disruption, i.e. tape stripping, was used. By repeatedly pressing an adhesive film to the surface of the skin and then abruptly removing it, only the uppermost skin layers, the stratum corneum, gets abrased. Unlike laser microporation, this technique has to be rated as much less invasive. Also the tape-stripped skin area is inevitably larger compared to the constrained area of 10 mm^2^ treated with the laser, on which the (epi)dermal tissue between the micropores is left intact [16]. In the above mentioned work, communication between epithelial cells and DETCs via the activating receptor NKG2D was crucial for induction of TH2-biased responses. One could speculate that repeated tape-stripping induces a more pronounced upregulation of the NKG2D ligand Rae-1 on keratinocytes than laser microporation, thereby leading to enhanced activation of DETCs. Finally, besides the diverging technique used for barrier disruption, Strid et al. applied not only a different allergen, i.e. ovalbumin, but also a higher dosage (100 µg vs. 1 µg) compared to our approach, a fact that could also contribute to the discrepant outcomes.

Taken together, our data indicate that systemic immune responses following laser-mediated EPI are not significantly modulated or regulated by γδ T cells, however; we propose that γδ T cells are rather responsible for shaping the local inflammatory milieu under steady state conditions.

MCs act as potent producers of various pro-inflammatory and TH2 cytokines [40, 41] and have been attributed to play a role in DC stimulation and migration. Furthermore, MCs can both directly or indirectly regulate T cell recruitment, activation, proliferation, and cytokine secretion (reviewed in [42]). Therefore, we hypothesized that MC and basophil depletion would impair TH2 polarization and consequently promote TH1 responses.

However, the cytokine and chemokine milieu in the skin after laser microporation was largely unaffected by the depletion of MCs with the exception of TSLP and GROα, which were significantly downregulated in MC-depleted mice. Similarly, a decreased induction of MIP-1α, and MIP-2α was detected in the absence of MCs (not significant). Interestingly, we observed that DT treatment by itself caused the induction of most skin chemokines. We thus reasoned that the DT-induced depletion of MCs leads to the release of immune mediators and alarmins from necrotic MCs and causes the activation of other skin-resident innate cells.

Nevertheless, depletion of MCs had only slight effects on the adaptive T cell response after EPI. Consistent with our stated hypothesis, MC-depleted mice show significantly decreased Phl p 5-specific serum IgG1 titers compared to C57BL/6 control mice after EPI (Fig. 2A). Thus, it should be considered that MC play a potential role in the TH2-dependent class switch to IgG1. In contrast to those findings, antigen-specific cell-bound IgE levels were not affected by the presence of MCs (Fig. 2B). However, we found no difference in the secretion of pro-inflammatory cytokines by antigen-restimulated lymphocytes between MC-depleted and WT C57BL/6 control mice (Fig. 3B). Unexpectedly, we observed an increased production of IL-4 and IL-10 by antigen-restimulated splenocytes in MC-depleted mice contrasting our finding of reduced serum IgG1 (Fig. 2A). As basophils, which are known to secrete large amounts of IL-4 after IL-33-stimulation, are depleted alongside MCs in our model, this is even more surprising. Thus, our results are contrary to data proposing either a role of MCs in promoting TH2 immunity in response to skin barrier disruption [43] or in promoting TH1/TH17 polarization [44]. Since cytokine expression was measured 12 days after the second immunization has been performed, we speculate that MCs and basophils only influence the early cytokine expression, but are rather dispensable in shaping the adaptive immune response after two or more epicutaneous immunizations using laser microporation.

FACS analysis revealed that MC-depleted mice show a significantly higher percentage of naïve T cells among isolated lymphocytes, compared to WT control mice (Fig. 4) and a concomitant decrease of effector T cells, confirming a role of MCs in T cell activation/maturation. This is consistent with data indicating a role for MCs in priming of DCs via ICAM/LFA-1 that can in turn enhance T cell priming [29, 45]. In line with our cytokine data, we observed no difference in transcription factor expression between MC-depleted mice and WT mice, thereby suggesting that MCs are dispensable for T helper cell polarization via transcription factor expression. Collectively, we propose that MCs are mostly dispensable for humoral TH2-immune responses in laser-mediated EPI, but may play a role in T cell activation/maturation. Finally, we concluded that MCs contribute to the laser-induced cytokine profile by either secretion or regulation of TSLP and GROα.

## 6. Conclusion

In summary, we did not observe a major role of γδ T cells or MCs in shaping the immune response after laser-mediated EPI with Phl p 5 mannan. However, they can contribute to the steady-state or damage-induced cytokine milieu in the skin, respectively.

## 7. Limitations of the study

C57BL/6 mice have the same genetic background as Mas-TRECK mice and were furthermore born and held in the same animal facility. While they should be a suitable control group in respect of certain factors like their specific microbiome, they did not undergo DT treatment and are therefore not an optimal control. Due to their low expression of the human DTR, DT treatment also results in transient basophil depletion. Furthermore, we have shown that DT-treatment can alter the chemokine milieu in the skin. Thus, immunological outcomes in the subsequent experiments cannot be considered solely a MC-dependent effect, but rather have to be considered MC- or basophil-dependent or DT-induced. This is of special interest regarding TH2 cytokine expression, since basophils are known to be producers of IL-4 and hence able to promote TH2 polarization. To overcome these limitations and exclusively elucidate the role of MCs in EPI in further studies novel constitutive or inducible MC-deficient models like Mcpt5-Cre, R-DTA, Mcpt5-Cre or iDTR might be empIoyed.

## 8. Acknowledgements

The authors would like to thank Dr. Masato Kubo, and Dr. Marcus Maurer for providing Mas-TRECK mice for this study.

