## Supplementary Data for "Mast cells and γδ T cells are largely dispensable for adaptive immune responses after laser-mediated epicutaneous immunization"

1.1 **Methods**

***Generation of allergen-mannan glycoconjugates***

Allergen-mannan neoglycoconjugates were generated by mild oxidation of mannan and subsequent reductive amination. For this purpose, 50mg of mannan (α-1,6-glycosidic-linked mannan from *S. cerevisiae*, Sigma) was dissolved in 1 mL of NaIO_4_ solution (11.91 mg/mL) and incubated at RT for 1 h under constant stirring in the dark, thereby resulting in an oxidation degree of 20%. For removal of excess NaIO_4_, a NAP-5 column (prepacked with sephadex G-25, GE Healthcare) was used according to the manufacturer’s instructions. For coupling of oxidized mannan to Phl p 5, mannan was mixed with recombinant Phl p 5 (LPS content <0.3 pg/µg) to reach a ratio (w/w) of 5:1 between mannan and the allergen. 12.5% of freshly prepared NABH_3_CN (sodium cyanoborohydride; 20 mg/mL in dH_2_O) was added and the reaction was incubated at RT under constant stirring in the dark overnight. To reduce any remaining aldehyde groups, 10% of NaBH_4_ (5 mg/mL in 50 mM borate buffer, pH 9.5) was added to the reaction, followed by a 6 h incubation at 4°C under constant stirring in the dark.

***Analysis and characterization of neoglycoconjugates***

The newly generated Phl p 5 mannan glycoconjugates were subjected to size exclusion chromatography using a 16/60 Sephacryl S-300 HR column (fractionation range: 10.000-1.500.000 Da, GE Healthcare) with an ÄKTA chromatographic system (GE Healthcare). The coupling efficiency of the neoglycoconjugate fractions was monitored by applying the fractions to a 12.5% reducing SDS-PAGE using colloidal Coomassie staining for protein visualization and fractions containing high molecular weight conjugates were pooled. The hydrodynamic radius of this glycoconjugate pool was measured using dynamic light scattering. Measurements were performed at a protein concentration of 1 mg/mL in DPBS at 40% laser power for 20x5 s on a dynamic light scattering instrument (DLS 802, Viscotek).

***Toluidine blue staining (mast cell depletion)***

To confirm the absence of mast cells at the time of immunization, small samples of ear tissue from Mas-TRECK mice were excised 18 days after the last treatment with diphtheria toxin (DT). Therefore, mice were put under light isoflurane (2.5% v/v) anaesthesia and lidocaine cream was applied. Tissue samples were embedded in O.C.T compound (Tissue-Tek, Sakura) and frozen in the gaseous phase of a liquid nitrogen chamber. 5 µm cryo-sections were generated using a cryostat microtome (Leica CM1950) and mounted on poly-L-lysine coated glass slides (Thermo Scientific). Slides were washed in PBS-T (PBS with 0.05% Tween^®^-20) before they were stained by dipping the slides in a 50 mL Greiner tube of toluidine blue working solution (2.8 g toluidine blue in a solution of 4 g urea dissolved in 120 mL dH_2_O and 280 mL of 70% isopropanol) for ten times. Slides were washed with PBS-T before they were destained by dipping into 50 mL of 70% isopropanol with 0.5% HCl. After destaining, sections were immediately analysed under a light microscope (Hun, Wilovert 30).

***Flow cytometry (basophil depletion)***

After 250 ng diphtheria toxin (DT) was intraperitoneally injected for five consecutive days, blood samples were drawn from the saphenous vein to confirm depletion of basophils in Mas-TRECK mice. Blood samples were diluted (1:10) with 1.5 mg/mL Li-heparin in PBS to avoid coagulation. An equal volume of staining mix was added containing anti-mouse CD3 (APC-labelled; clone 145-2C11, BioLegend), anti-mouse CD45-B220 (PerCp-Cy5-labelled; clone RA3-6B2, BioLegend), and anti-mouse IgE (FITC-labelled; clone RTK2071, BioLegend) diluted 1:50 in FACS buffer (1% BSA, 2 mM EDTA in PBS). Samples were incubated for 30 min at 4°C in the dark. After incubation, blood samples were washed with 400 µL FACS buffer before erythrocyte lysis was performed by addition of 300 µL 1xRBC lysis buffer (eBioscience). After 5 min of incubation at RT, cells were washed with 600 µL of FACS buffer. The supernatant was discarded and the cell pellet was dissolved in 100 µL FACS buffer. The samples were then analysed on a FACS Canto II (BD Biosciences) by gating for basophils (anti-IgE^+^, anti-CD45-B220^-^, and anti-CD3^-^).

***Epidermal sheets (monitoring of γδ T cells)***

To determine the presence of γδ T cells in the epidermal layer of the skin, epidermal sheets were generated from ear tissue after mice were sacrificed by cervical disclocation. Therefore, 3.8% NH_4_SCN (ammonium thiocyanate in dH_2_O; Carl Roth GmbH) was warmed to 37°C before a 500 µL drop was placed into a petri dish (35 mm). The dorsal and ventral side of the ear were separated and incubated on the drop of 3.8% NH_4_SCN with the inner side facing down for 20 min. The dorsal and ventral side of the ear were washed on a drop of PBS in a 24-well plate (Greiner) before the epidermis was separated from the dermis. The separated epidermis was fixed by floating on acetone for 1-2 min at RT, before it was washed again by floating on a drop of PBS. The epidermal sheet was then placed on a drop of staining mix containing anti-mouse CD3 (AlexaFluor647-labelled; clone 172A, BioLegend) diluted 1:200 in PBS and incubated for 30 min at 37°C. Nuclei were stained by putting the epidermal sheet on a drop of PBS containing Hoechst 33342 staining dye (dilution 1:1000; Thermo Fisher) for 3-5 min at RT. The epidermal sheets were washed twice with PBS and once with dH_2_O before they were mounted on glass slides using Roti Mount Aqua (Carl Roth GmbH). To detect γδ T cells, the slides were placed under a fluorescence microscope (Olympus IX70) with an appropriate filter.

***Generation of skin micropores and immunization***

One day prior to laserporation, an area of roughly 3 cm^2^ at the lower back of mice was shaved using an electric clipper, followed by treatment with depilatory cream (Veet sensitive). The depilatory cream was applied for 30 s under light anaesthesia (2.5% v/v isoflurane) before it was removed with a wet sponge. The next day, mice received ketamine/xylazine anaesthesia (80 mg ketamine and 7.5 mg xylazine per kg body weight) by intraperitoneal injection. Mice underwent laser microporation and the allergen suspension (1 µg of Phl p 5 mannan in 20 µL DPBS) was applied onto the laser-induced micropores using a pipette tip. After all the liquid was taken up, the treated area was covered with an adhesive tape (OPSITE Flexifix, Smith & Nephew).

***IgG1 ELISA***

A luminometric ELISA was performed to determine the production of Phl p 5-specific serum IgG1 after immunization with Phl p 5 mannan glycoconjugates. Therefore, a 96-well plate (flat white chimney; Greiner) was coated with 1 µg/mL recombinant Phl p 5 diluted in PBS (50 µL per well) overnight at 4°C. After incubation at RT for 1 h, the coating solution was discarded. 200 µL of blocking buffer (2% skim milk, blotting grade, and 0.1% Tween®-20 in PBS) was added to each well and incubated 1 h at RT. The plate was washed with PBS-T (PBS with 0.1% Tween-20) using an automated plate washer (Tecan 96PW Microplate Washer). Meanwhile, serum dilutions (1:1000 and 1:10000) were prepared in blocking buffer. 50 µL of each serum dilution was added to the wells and incubated for 1 h at RT. After another washing step, 50 µL of the secondary antibody (goat anti-mouse IgG1-HRP, Bio-Rad) was added as a 1:1000 dilution in blocking buffer. The plate was incubated for 1 h at RT. After a final washing step, 50 µL of an ELISA BM chemiluminescence substrate (Roche) was added and after 2-3 min the luminescence was measured in relative light units (RLU) using a plate reader (Tecan inifinite 200Pro; integration time of 1000 ms and attenuation set to automatic).

***Basophil activation test***

For the measurement of cell-bound allergen-specific IgE levels, a basophil activation test was performed. Briefly, 30 µL of heparinized blood sample was mixed with either 30 µL RPMI (stimulation control), 30 µL RPMI with 20 ng/mL rPhl p 5 (final concentration 10 ng/mL) or 30 µL RPMI with 200 ng/mL rPhl p 5 (final concentration 100 ng/mL) in a 96-well plate (V-bottom plate; Greiner) and incubated at 37°C and 5% CO2 for 2 h. 80 µL of ice-cold FACS buffer (1% BSA, 2mM EDTA in PBS) was added to each well and the plate was spun down at 260 g for 5 min at 4°C. The supernatants were aspirated and pellets were resuspended in 30 µL staining mix containing anti-mouse IgE (FITC-labelled; clone RME-1, BioLegend), anti-mouse CD4 (PerCp/Cy5.5-labelled; clone GK1-5, BioLegend), anti-mouse CD200R (APC-labelled; clone OX110, eBioscience), and anti-mouse CD19 (PE/Cy7-labelled; clone eBio1D3, BD Biosciences), all diluted 1:200 in FACS buffer and incubated on ice for 25 min. Then 80 µL of FACS buffer was added and the plate was spun down for 5 min at 260 g at 4°C. The supernatants were aspirated and the pellet was resuspended in 100 µL 1xRBC lysis buffer (eBioscience). After 5 min of incubation at RT, the plate was spun down, supernatants were discarded and 100 µL FACS buffer was added. After a final washing step, the pellets were dissolved in 50 µL FACS buffer and the median fluorescence intensity of CD200R was analysed on a FACS Canto II flow cytometer (BD Biosciences, Franklin Lakes, New Jersey) by gating on non-T/non-B/IgE^high^ cells.

*Gating strategy for the basophil activation test:*

FSC/SSC were used to gate for the cell population and the FSC height and width were used to determine singlet cells. Antibodies specific for the extracellular markers CD4 and CD19 were used to exclude T cells and B cells, respectively. Therefore, to gate on the basophil population IgE^+^, CD4^-^ and CD19^-^ cells were considered. Activation status of the basophil population following the treatment was determined by analysis of the median fluorescence intensity of CD200R.

***T helper cell polarization***

After mice were sacrificed, spleens and skin-draining lymph nodes (inguinal, axillary, and brachial) were prepared. Lymph nodes and spleens were homogenized and after sedimentation of debris, the monodisperse supernatants of LN samples were transferred into 15 mL tubes with DPBS. Red blood cell lysis was performed with spleen samples using Ammonium-Chloride-Potassium lysis buffer (ACK buffer; 0.15 M NH_4_Cl, 10 mM KHCO_3_, 0.1 mM Na_2_EDTA, pH 7.2). All samples were spun down at 260 g for 5 min. The supernatants were discarded and the pellets were washed with 5 mL DPBS. Pellets were dissolved in 1 mL (spleen) or 200 µL (lymph nodes) T cell medium (RPMI-1640; 10% FCS, 25 mM HEPES, 2 mM L-Glu, 100 µg/mL streptomycin, 100 U/mL penicillin). Cells were diluted to a concentration of 4x10^6^ cells/mL and 75 µL of each cell suspension was seeded in a sterile 96-well tissue culture plate (U-bottom; Greiner). 75 µL of Phl p 5 (10 µg/ml) was added and the plate was incubated at 37°C and 5% CO_2_. After 5 days of incubation, culture supernatants were removed and the remaining restimulated lymphocytes were washed twice with 100 µL of DPBS before 20 µL of supernatant from anti-CD16/32 antibody (clone 2.4G2) producing hybridoma (ATCC number HB-197) was added to the cells to block FcγR and incubated for 5min at 4°C in the dark. 20 µL of extracellular staining mix were added to each well containing a live/dead staining (fixable viability dye eFluor 506; Thermo Fisher; 1:1000), anti-mouse CD62L (FITC-labelled; clone MEL-14, BioLegend; 1:200), anti-mouse CD44 (BV650-labelled; clone IM7, BioLegend; 1:100), and anti-mouse CD4 (APC-Cy7-labelled; clone GK1.5, eBioscience; 1:400) in DPBS. After 30 min incubation at 4°C in the dark, 100 µL of ice-cold FACS buffer was added to each well. The cells were spun down and the supernatant was discarded. The cells were then resuspended in 150 µL ice-cold Fix/Perm buffer (FoxP3 staining buffer kit; eBioscience). The cells were incubated for 60 min at 4°C in the dark before they were spun down and the supernatant was discarded. Two washing steps with 100 µL Perm buffer (FoxP3 staining buffer kit; eBioscience) were performed before the cell pellets were dissolved in 20 µL of 10% naïve mouse serum in Perm buffer for additional blocking. After 5 min of incubation at RT, 20 µL of the intracellular staining mix was added containing anti-mouse FoxP3 (APC-labelled; clone FJK-16s, Invitrogen; 1:100), anti-mouse T-bet (PE-Cy7-labelled; clone 4B10, BioLegend; 1:100), anti-human/mouse GATA3 (BV421-labelled; clone 16E10 A23, BioLegend; 1:200), and anti-mouse RORγT (PE-labelled; clone B2D, Invitrogen; 1:100). Cells were incubated with mouse serum and intracellular staining mix at RT in the dark for 30 min followed by two final washing steps with 100 µL Perm buffer were performed. Cells were resuspended in 80 µL FACS buffer (1% BSA, 2 mM EDTA in PBS) and analysed using a CytoFLEX S flow cytometer (plate mode; Beckman Coulter).

*Gating strategy:*

FSC/SSC were used to gate for the lymphocyte population and the FSC height and width were used to determine singlet cells. Antibodies specific for the extracellular markers CD4, CD62L (lymph node homing) and CD44 as well as a live/dead viability dye were used to differentiate between naïve, central memory, and effector CD4^+^ live lymphocytes.

***Cytokine production***

After isolated lymphocytes from spleen and lymph nodes were restimulated *in vitro* with 10 µg/mL Phl p 5 for five days, supernatants were transferred into a new 96-well plate (flat bottom; Greiner). For quantification of the cytokines TNF-α, IFN-γ, IL-4, IL-5, IL-6, IL-10, IL-13, IL-17, and MCP-1 produced by *ex vivo* restimulated lymphocytes from immunized TCRd mice, ProcartaPlex multiplex cytokine panels (Invitrogen) were used according to the manufacturer’s protocol. For quantification of cytokines TNF-α, IFN-γ, IL-2, IL-4, IL-5, IL-6, IL-10, IL-13, IL-17A, IL-17F, IL-21, IL-22 produced by *ex vivo* restimulated lymphocytes from immunized mast cell-depleted Mas-TRECK mice and C57BL/6 mice a LegendPlex Mouse TH cytokine panel (BioLegend) was performed according to the manufacturer’s protocol.

***Laser-induced cytokine-milieu in murine skin***

Mice were shaved and depilated one day prior to laser treatment as described for immunizations. Two non-overlapping areas of 1 cm^2^ on the back were microporated using the P.L.E.A.S.E laser device (laser settings: 8.3 J/cm^2^, 50 µs pulse duration, 500 Hz frequency, 5% pore density, 3 pulses). 6 h and 24 h after laser microporation the mice were sacrificed by cervical dislocation. Control groups that were not microporated were both shaved and depilated after they had been sacrificed. The lasered skin area (non-microporated skin in the case of the control group) was excised and transferred into 12 mL tubes. The weight of the skin tissue was determined and a 20x amount of DPBS (+1% protease inhibitor cocktail; Sigma, P8340) was added. The skin was homogenized on ice using an IKA® ULTRA-TURRAX® tissue homogenizer (speed of 18000/min; 6x10 s, 10 s breaks in between). The homogenized skin samples were transferred into 1.5mL Eppendorf tubes and centrifuged at >20.000 g for 15 min. The supernatants were filtered through a 0.22 µm SpinX column (Corning) before the concentrations of MIP1α, RANTES, GROα, MIP2α, MCP1, MCP5, MIP3α, IP-10, IL-6, TNF-α, and TSLP (for TCRd mice), and TSLP, IL-6, TNF-α, MCP1, MIP1α, RANTES, MCP3, IP-10, GROα, and MIP2α (for Mas-TRECK mice) were analysed using ProcartaPlex multiplex assay (eBioscience) as described before.

1.2 **Results**

***Purification and characterization of Phl p 5 mannan neoglycoconjugates***

*Size exclusion chromatography*

Size exclusion chromatography (SEC) was performed to separate the Phl p 5 mannan neoglycoconjugates by size. Therefore, a HiPrep 16/60 Sephacryl S-300 HR column was used on an ÄKTA prime chromatography system. 2 mL of the freshly generated Phl p 5 mannan neoglycoconjugates were loaded onto the sample loop with a constant flow rate of 0.5 mL/min. Elution was performed with PBS and the eluate was collected in 3 mL fractions starting at minute 0 of the SEC until after the first peak ended at minute 200. Due to the larger size, mannan conjugated Phl p 5 is eluted first as its size prevents retention in the pores of the adsorbent. Therefore, the first visible peak in the chromatogram (Suppl. Fig. 1) represents the eluted neoglycoconjugates (starting at minute 60 of SEC), whereas the second peak (starting at minute 200 of SEC) resembles unconjugated mannan that also absorbs at 280 nm.

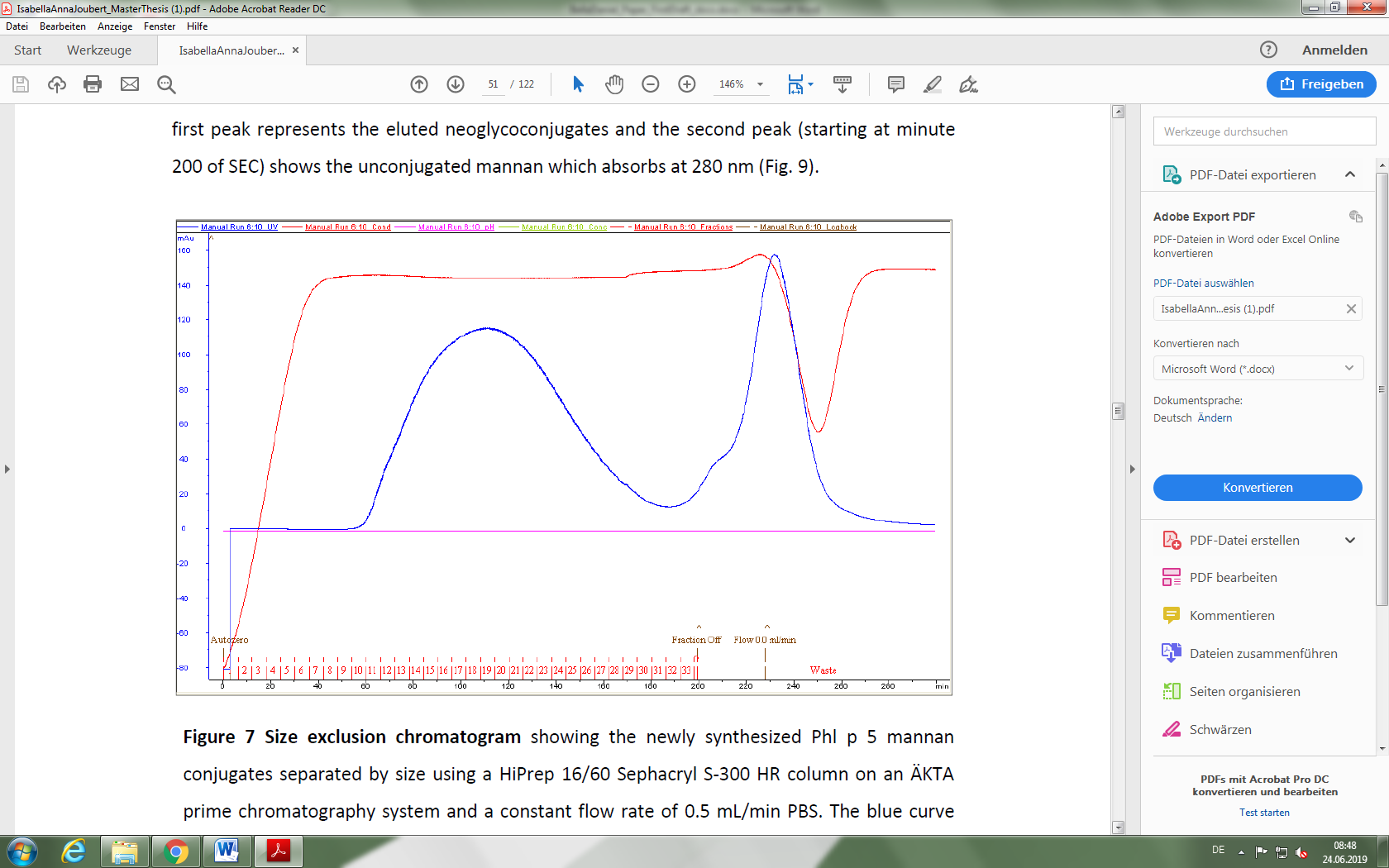

**Supplementary Figure 1. Size exclusion chromatogram of P5-MN neoglycoconjugates.** P5-MN conjugates were separated by size using a HiPrep 16/60 Sephacryl S-300 HR column on an ÄKTA prime chromatography system. The blue and the red curve represent changes in the absorbance at 280 nm and in conductivity/absorbance, respectively.

***Reductive amination is an effective way to generate allergen-neoglycoconjugates***

After collecting the different fractions, we wanted to investigate whether the separation by SEC was efficient and resulted in the enrichment of high molecular weight Phl p 5 mannan neoglycoconjugates. To evaluate molecular weight (MW) and also coupling efficiency of the individual fractions after SEC, we performed an SDS-PAGE on a reducing 12.75% gel and visualized proteins by staining with colloidal Coomassie. Suppl. Fig. 2 clearly depicts that the MW of the individually obtained fractions (11-21) was approximately higher than 100 kDa, i.e., more than 3-fold larger compared to unconjugated Phl p 5 (MW = 30kDa). In addition, our gel images revealed no detectable unconjugated Phl p 5 in all analysed fractions (1-33). Thus, SDS-PAGE verified the effective and controlled generation of high molecular weight Phl p 5 mannan neoglycoconjugates with a high coupling efficiency.

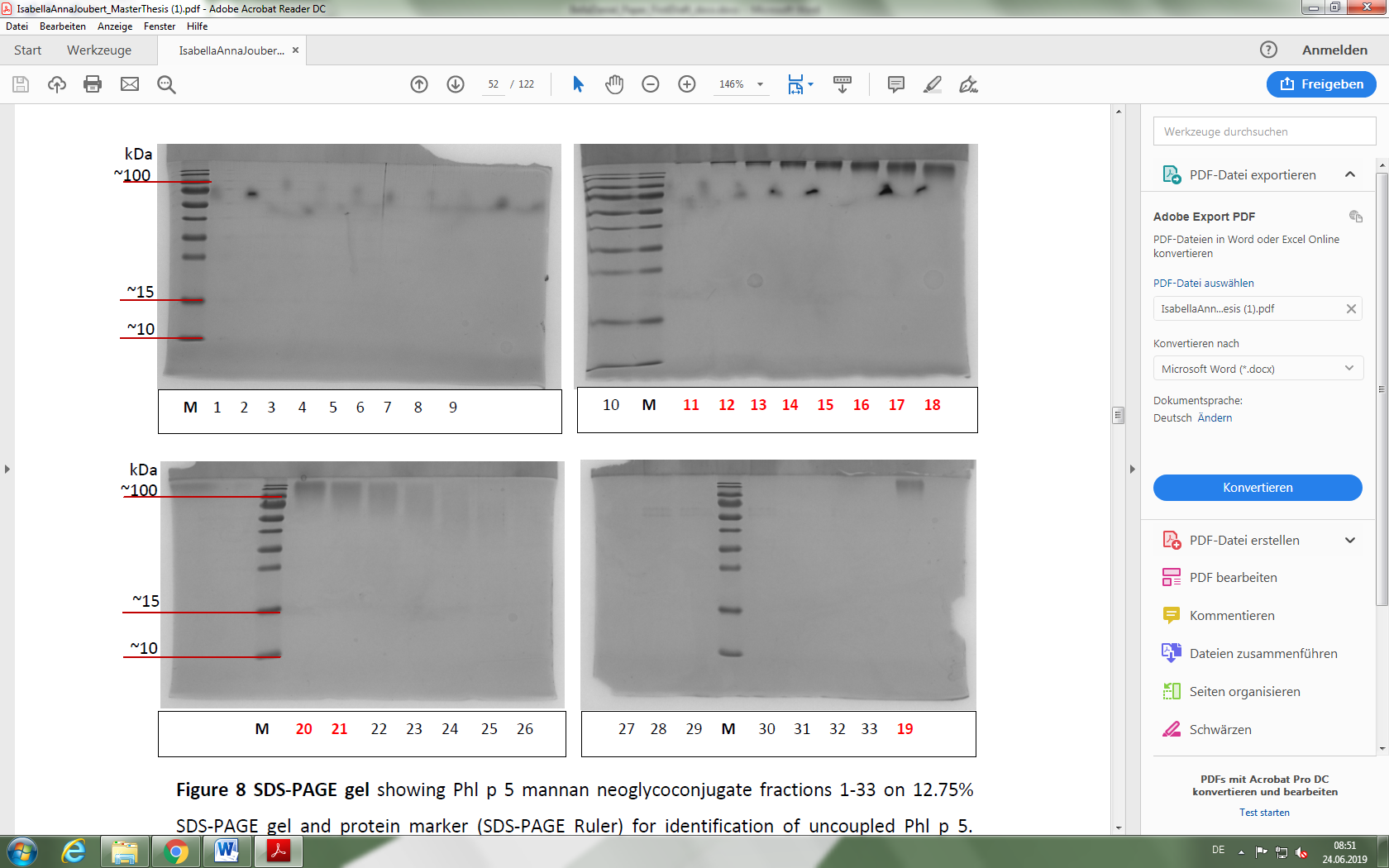

**Supplementary Figure 2. SDS-PAGE of P5-MN neoglycoconjugate fractions.** Fractions 1-33 were analysed on a 12.75% gel. PageRuler™ Prestained Protein Ladder, 10 to 180 kDa was used as marker for identification of uncoupled Phl p 5. Fractions marked in red were pooled for further analysis.

***Phl p 5 mannan neoglycoconjugates are heterogeneous in size***

To determine the hydrodynamic radius and characterize size distribution profile of the Phl p 5 mannan glycoconjugates, we performed a dynamic light scattering (DLS) analysis of the pooled fractions 11-13 in PBS. Pooled fractions were repeatedly measured at a protein concentration of 1 mg/mL using the solvent settings for PBS. As expected, the mean hydrodynamic radius of the neoglycoconjugate pool estimates to 10.1 nm (Suppl. Fig. 3) and therefore is 3-fold higher than the mean hydrodynamic radius of unconjugated Phl p 5 (R_H_ = 3 nm, not shown). Furthermore, the obtained size distribution demonstrates heterogeneity, suggesting the presence of bigger aggregates in the solution.

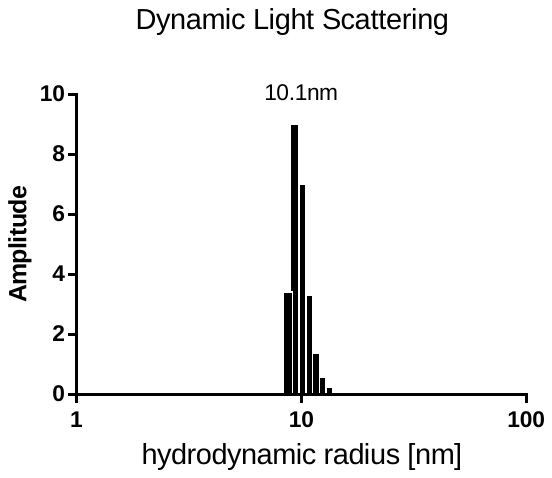

**Supplementary Figure 3. Measurement of the hydrodynamic radius and size distribution of pooled fractions 11-13 by DLS.**

***Visual verification of γδ cells and mast cells in skin samples***

Epidermal sheets were prepared from the ears from sacrificed mice at the end of the immunization experiments to verify the presence and absence of γδ T cells in TCRd^-/+^ and TCRd^-/-^ mice, respectively. Suppl. Fig. 4 shows representative examples of epidermal sheets from TCRd^-/+^ (A, B) and TCRd^-/-^ (C, D) mice. Consistent with their genotype, epidermal sheets from TCRd^-/-^ showed no detectable γδ T cells, whereas in TCRd^-/+^ γδ T cells with a dendritic morphology were observed. Thus, we were able to validate TCRd^-/-^ mice as a suitable model to study effects depending on the presence of γδ T cells.

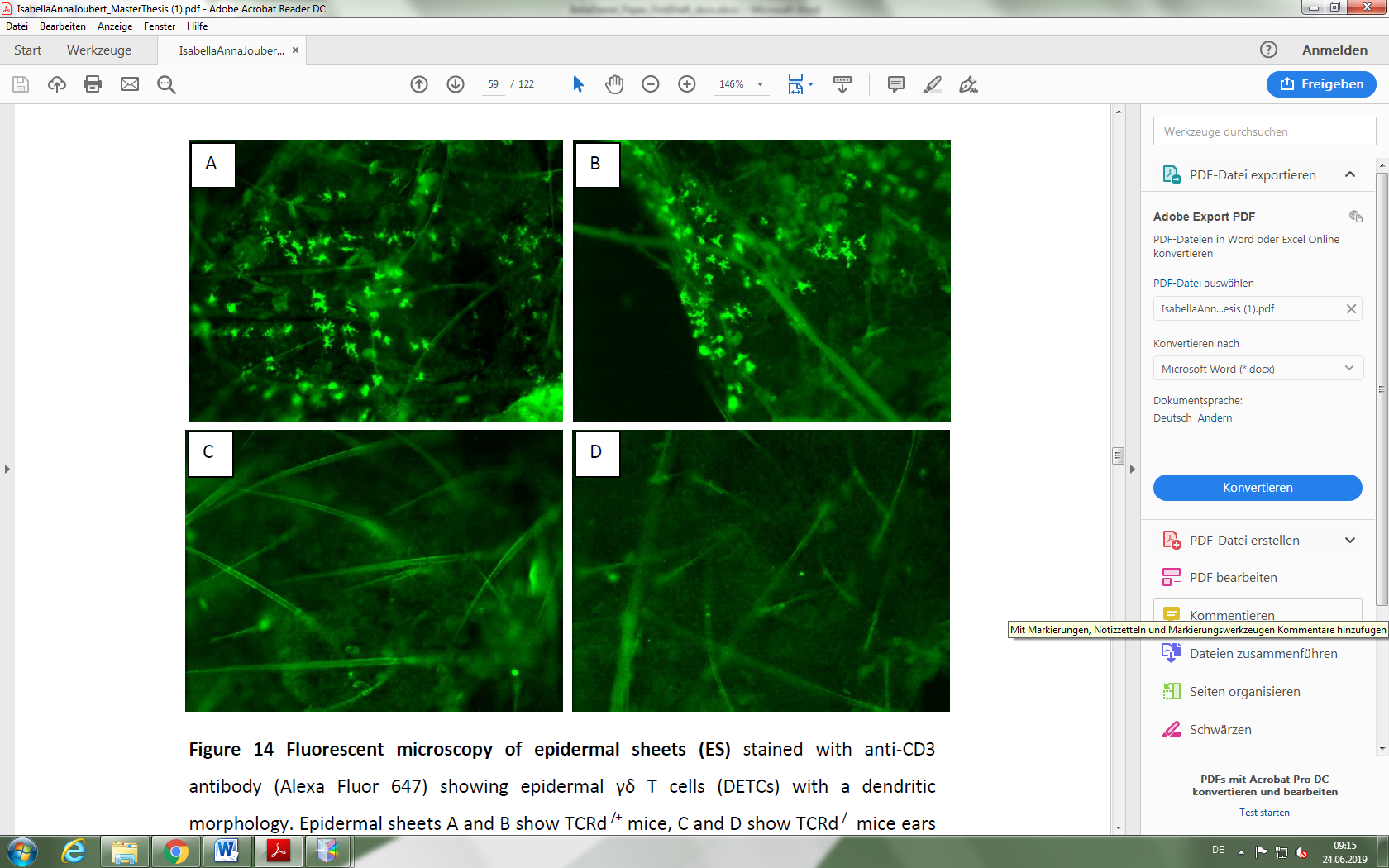

**Supplementary Figure 4. Fluorescence microscopy pictures of epidermal sheets.** Epidermal sheets (ES) from TCRd^-/+^ mice (A and B) or TCRd^-/-^ mice (C and D) were stained with anti-CD3 antibody (Alexa Fluor 647) to visualize γδ T cells in the epidermis.

To demonstrate that MCs were depleted at the time of immunization, small samples of ear tissue were prepared after the first immunization (day 18) followed by a histological analysis. It has been reported that MCs and basophils are depleted for at least 12 days after cessation of DT treatment [28], and we verified that MCs were still depleted on day 18 after the last DT application. Cryo-sections of both MC-depleted Mas-TRECK mice and C57BL/6 control mice were prepared and metachromatically stained with toluidine blue to visualize connective-tissue MCs in the skin. We only observed MCs in C57BL/6 control mice (Suppl. Fig. 5; C and D) as identified by their red-purple color. In contrast, DT-treated Mas-TRECK cryo-sections (Suppl. Fig. 5; A and B) displayed a complete absence of MCs. Surprisingly, MC were still completely absent from the connective-tissue in the skin at day 18. Taken together, we were able to confirm the conditional, temporal and specific depletion of MCs in skin of Mas-TRECK mice by application of DT.

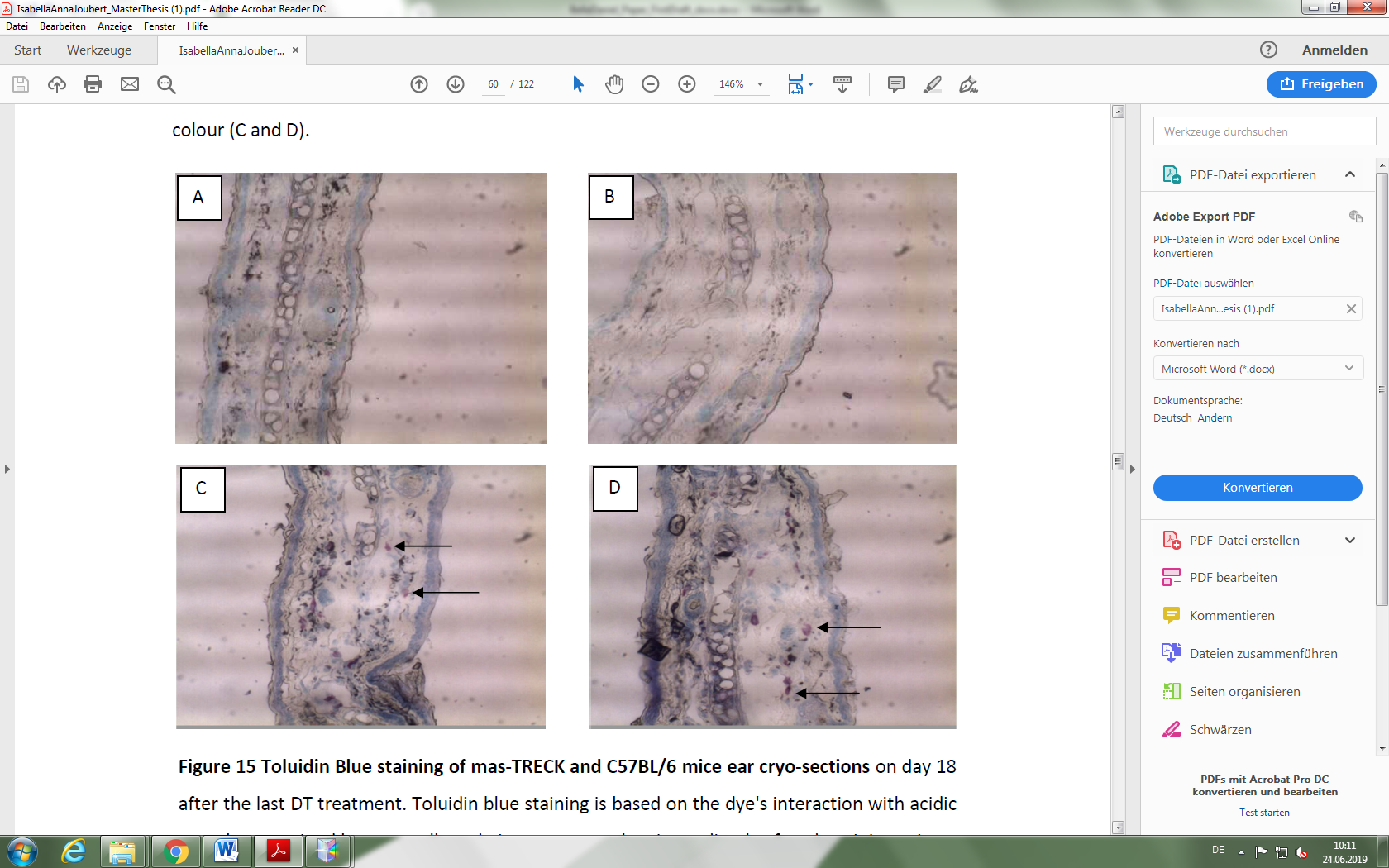

**Supplementary Figure 5. Microscopic pictures of cryo-sections from ear skin.** To visualize connective-tissue MCs in the skin of Mas-TRECK mice (A and B) and C57BL/6 controls (C and D), cryo-sections from ear skin were prepared on day 18 after the final DT treatment and metachromatically stained with toluidine blue.

***Antigen-restimulated lymphocytes secrete elevated levels of TH2 cytokines in the absence of MCs***

Isolated lymphocytes from skin draining lymph nodes from immunized Mas-TRECK and C57BL/6 mice were restimulated with 10 µg/mL recombinant Phl p 5 *in vitro* for five days and cytokine secretion in the culture supernatants was measured by LegendPlex multiplex analysis. We observed that lymphocytes isolated from MC-depleted mice show a consistent, however non-significant, trend of higher TH2 cytokine secretion compared to C57BL/6 control mice (Suppl. Fig. 6). This data points towards a role of MCs in polarization T helper type 2 cells.

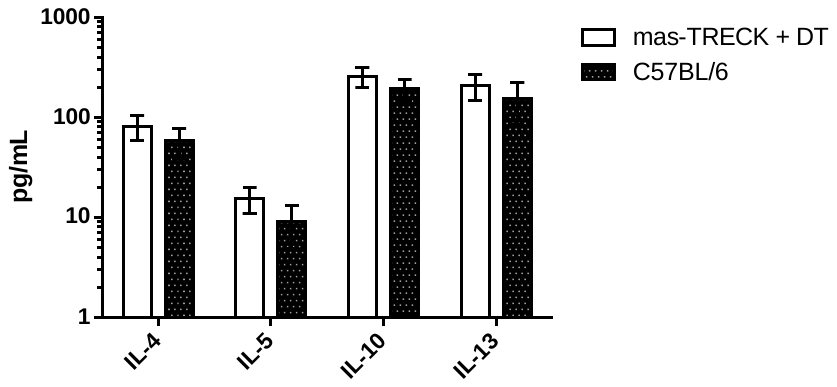

**Supplementary Figure 6. Impact of MC depletion on TH2 cytokine secretion by antigen-restimulated lymphocytes.** Skin-draining LN cells were isolated after EPI with P5-MN and cultured for 5 days in the presence of Phl p 5. Datasets for DT-treated Mas-TRECK (n=10) and C57BL/6 (n=6) mouse groups were analysed using a one-way ANOVA and Tukey’s post hoc test with a single pooled variance and a significance level of 95% using GraphPad Prism 6 software. Data is shown as means ± SEM.
